## Supplementary Table 1 for "USP8 inhibition promotes Parkin-independent mitophagy in the *Drosophila* brain and in human neurons"

KEGG PATHWAY

| Index | Name | p-value | Adjusted p-value | Z-score | Combined score |
| --- | --- | --- | --- | --- | --- |
| 1 | Homologous recombination | 0.002971 | 0.03743 | -74.68 | 434.55 |
| 2 | Dorso-ventral axis formation | 0.0004092 | 0.008594 | -47.90 | 373.66 |
| 3 | Hedgehog signaling pathway | 0.01231 | 0.07754 | -61.20 | 269.11 |
| 4 | Glycolysis / Gluconeogenesis | 0.03123 | 0.1474 | -74.52 | 258.30 |
| 5 | Glycerolipid metabolism | 0.09940 | 0.2853 | -70.00 | 161.60 |
| 6 | AGE-RAGE signaling pathway in diabetic complications | 0.05870 | 0.1947 | -56.70 | 160.75 |
| 7 | FoxO signaling pathway | 0.009388 | 0.07393 | -33.05 | 154.28 |
| 8 | Phototransduction | 0.000007766 | 0.0004893 | -12.60 | 148.22 |
| 9 | Mitophagy | 0.07635 | 0.2405 | -50.05 | 128.75 |
| 10 | Fanconi anemia pathway | 0.03979 | 0.1474 | -30.98 | 99.90 |
